# On the energetics and stability of a minimal fish school

**DOI:** 10.1101/596023

**Authors:** Gen Li, Dmitry Kolomenskiy, Hao Liu, Benjamin Thiria, Ramiro Godoy-Diana

## Abstract

The physical basis for fish schooling is examined using three-dimensional numerical simulations of a pair of swimming fish, with kinematics and geometry obtained from experimental data. Energy expenditure and efficiency are evaluated using a cost of transport function, while the effect of schooling on the stability of each swimmer is examined by probing the lateral force and the lateral and longitudinal force fluctuations. We construct full maps of the aforementioned quantities as functions of the spatial pattern of the swimming fish pair and show that both energy expenditure and stability can be invoked as possible reasons for the swimming patterns and tail-beat synchronization observed in real fish. Our results suggest that high cost of transport zones should be avoided by the fish. Wake capture may be energetically unfavorable in the absence of kinematic adjustment. We hereby hypothesize that fish may restrain from wake capturing and, instead, adopt side-to-side configuration as a conservative strategy, when the conditions of wake energy harvesting are not satisfied. To maintain a stable school configuration, compromise between propulsive efficiency and stability, as well as between school members, ought to be considered.

## Introduction

The behaviors of living beings provide amazing examples of aggregated dynamics that result from complex social reasons [1–4]. Depending on the species, animals aggregate and modulate group cohesion to improve foraging and reproductive success, avoid predators or facilitate predation. Global cohesive decision and action for the whole group result from different types of interaction at the local scale. Fish schools, for instance, are an archetypal example of how local interactions lead to complex global decisions and motions [5]. Fish interact through vision but also by sensing the surrounding flow using their lateral line system [6]. From the fluid dynamics perspective, hydrodynamic interactions between neighbors have often been associated with swimming efficiency strategies, considering how each individual in the school is affected by the vortical flows produced by its neighbors. Breder [7] already recognized the importance of this issue, and more recent works have described how fish make use of vortices when swimming through an unsteady flow, whether produced by neighboring fish or by other features in the environment (see e.g. the review by Liao [8]). Concerning collaborative interactions between swimming fish, the first clear picture was proposed in the early 70’s by Weihs’ pioneering work [9]. He focused on interactions within a two-dimensional layer of a three-dimensional school, and proposed an idealized two-dimensional model in which each individual in the fish school places itself to benefit from the wakes generated by its two nearest neighbors, giving rise to a precise diamond-like pattern.

Weihs’ theory has been followed by extensive experimental verification generally comforting the idea of decreased energetic cost of locomotion in fish schools. Thus, Fields [10] reported decreased tail beat frequency as indicator to decreased swimming effort in groups of pacific mackerel (*Scomber japonicus*). Herskin and Steffensen [11] measured both tail beat frequency and oxygen consumption in sea bass *Dicentrarchus labrax*, and also found strong evidence for energy saving. Johansen et al. [12] estimated that trailing fish in a school (striped surfperch *Embiotoca lateralis*) benefited from over 25% reduction in oxygen consumption, based on correlations between swimming speeds, pectoral fin beat frequency, and oxygen consumption of solitary fish. Marras et al. [13] also inferred reduced costs of swimming from measurements of tail-beat frequency of grey mullet *Liza aurata* alone and in schools, combined with relationships between tail-beat frequency and activity metabolism. Interestingly, they found that all members of the school received energetic benefit regardless of their spatial position relative to neighbors. Halsey et al. [14] examined how water turbulence affected the tail beat frequency of sea bass *Dicentrarchus labrax* swimming in schools of different size. They reported a trend for attenuation of energy advantages which they explained by frequent short-term changes in fish position mediated by the turbulence. At the same time, they recognized that turbulence could modify the relationship between tail beat frequency and rate of oxygen consumption.

While consensus generally is maintained about energetic advantage of swimming in a group, the ubiquity of diamond formation as energy optimization policy has been subject to debate. Groups of red nose tetra fish *Hemigrammus bleheri* in shallow water, for instance, show strong preference for a phalanx configuration in high energy-demand swimming regimes [15]. Our understanding of the essential hydrodynamic interactions behind energy saving being insufficient to explain such behaviors observed in biological experiments, we resort in this work to the computational fluid dynamics (CFD) approach. Its most important advantage in the present context is that it provides a direct quantitative estimate to the hydrodynamic power in self-propelled swimming. Although the CFD modelling of collective swimming is not new, most of the prior work has been limited to groups of two-dimensional (2D) swimmers in 2D fluids [16–25].

We are only aware of two previous three-dimensional (3D) CFD studies of fish schooling. Large-eddy simulations by Daghooghi and Borazjani [26] modelled a large group of fish all swimming in the same plane as an infinite lattice of self-propelled in-phase synchronized swimmers. It was found that, for equal power, the fish in a rectangular formation with sufficiently small lateral distance swam 20% faster than alone. It was noticed, however, that the wake broke down into small, disorganized structures showing little evidence for constructive vortex interaction. As an alternative to the wake capture, channeling effect that enhances the flow velocity between swimmers was hypothesized to be the main energy-saving mechanism. A recent study by Verma et al. [27] included 2D and 3D numerical simulations. The 2D model was coupled with a deep reinforcement-learning algorithm to show that the collective energy savings in a fish school can be explained by a “smart” follower actively harvesting energy from the wake vortices behind its leader(s), achieving up to 32% increase in time-average swimming efficiency and 36% decrease in the cost of transport (*CoT*), with respect to a solitary swimmer. The control policy found in those 2D simulations was subsequently integrated within the 3D model in form of simplified rules. The 3D simulations showed 11% increase in efficiency and 5% decrease in *CoT*.

The topic of our present study is local hydrodynamic interaction between individuals in small schools of tetra fish, as described in earlier experimental work by Ashraf et al. [15, 28]. A physical description of the local interactions between nearest neighbors, which are crucial in determining the whole group dynamics, still needs deeper insight. We therefore study the minimal subsystem of fish school, consisting in two fish swimming together, using a three-dimensional computational approach developed by Li et al. [29–31]. We investigate the consequences of spatial organization and kinematic synchronization on the energy expenditure of the two-fish school (see Fig. 1) and the intensity of the pressure fluctuations induced by one individual on its neighbor. The fish are immersed in a sufficiently large numerical water channel (see Materials and Methods). In the following description, we call ‘protagonist fish’ the one for which we report the swimming performance data such as forces, power, etc. The other one is called ‘companion fish’. We prescribe the temporal deformation of the fish midline having the same functional form for both fish, but with a phase shift *δϕ* (positive when protagonist lags behind the companion). It is known from past experiments [15, 28] that groups of tetra fish maintain some particular fixed configurations and constant gaits (see, e.g., [32, Movie S1]). In our numerical study, we presume that all fixed configurations (i.e., fixed relative positions of the centers of mass (CoM) of the two fish) are realizable. We implement the simulations in order to clarify whether the observed configurations stand out in terms of favorable hydrodynamic interaction. Moreover, groups of tetra fish tend to align in one horizontal plane, i.e., the vertical offset between any two group members is smaller than each individual height [15, 28]. Considering that the hydrodynamic disturbances are the strongest in the same horizontal plane, we only investigate in-plane configurations in this work by imposing zero vertical separation between the two fish. The lateral spacing *δx* and the longitudinal spacing *δy* remain constant during each numerical simulation. Note that the protagonist is the follower and the companion fish is the leader if *δy <* 0, or vice versa if *δy >* 0. We perform a series of 312 simulations in total to realize parameter sweep in *δx*, *δy*. In addition, we test 4 different values of the phase shift *δϕ*. Fig. 2 shows a visualization of the three-dimensional flow in two typical swimming configurations.

**Fig 1.**
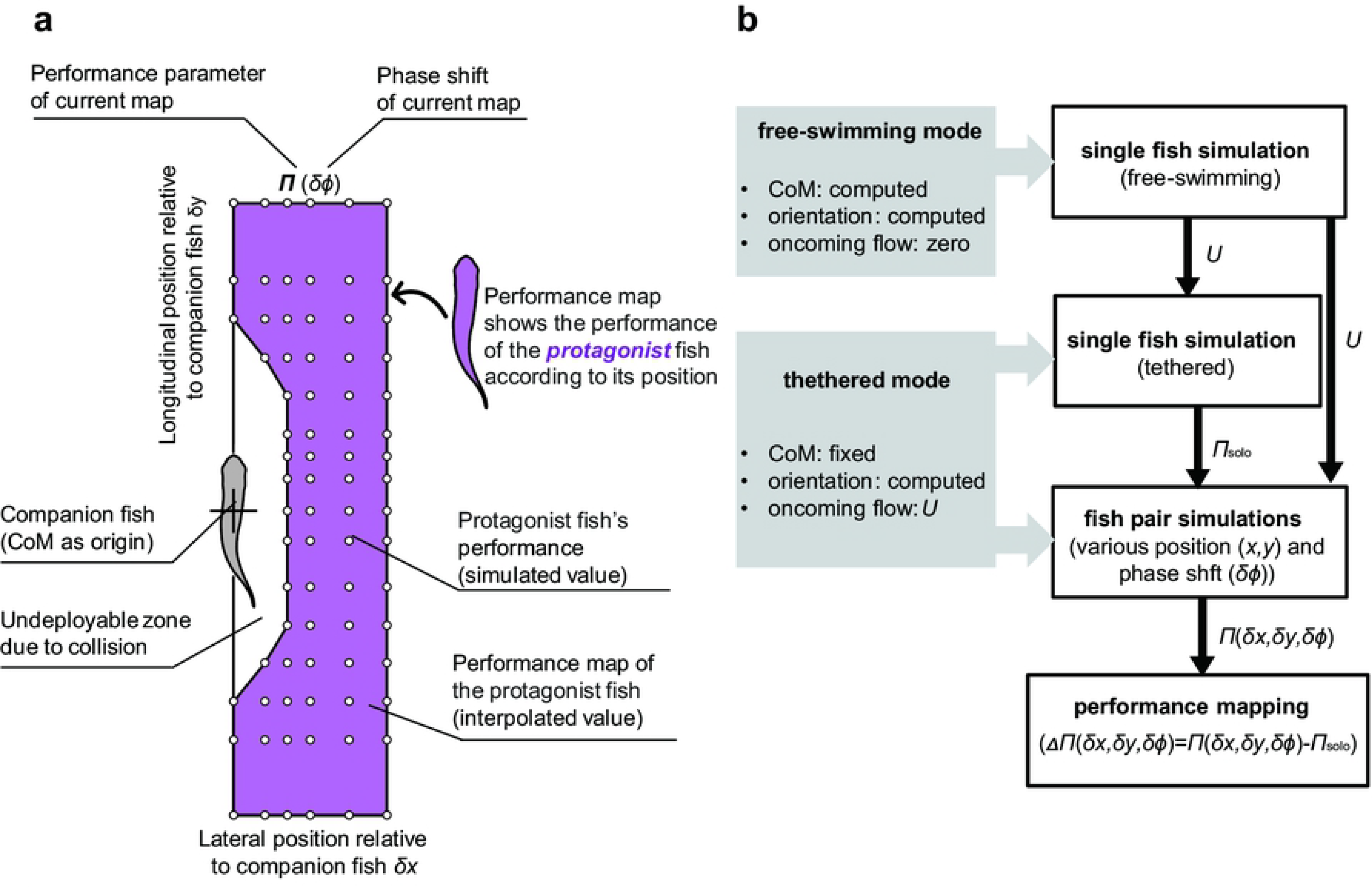
(a) An explanation diagram of performance maps in Figs. 3-Fig.6. Simulations were implemented by varying the relative longitudinal and lateral positions between two fish. To test the influence of phase difference, for each position (circles) we implemented four simulations (*δϕ* = 0, *T*/4, *T*/2 and 3*T*/4, respectively). Based on simulation results and interpolation, maps for swimming performance parameters were drawn. This performance map provide the performance value of the protagonist fish with its companion fish located at the origin. (b) (LHS) We conducted simulations in two modes: free-swimming (self-propelled) mode and tethered (fixed CoM) mode; (RHS) Procedure flow of simulations. Firstly, we simulated free-swimming single fish, obtained the terminal speed and apply to the rest simulations; We then simulated single fish swim and fish pair swim with CoM fixed. The relative performance of protagonist fish in fish pair to single fish, is used to draw a performance map to demonstrate the influence of relative position and phased shift comprehensively. Here *U* is the terminal speed in single fish free-swimming, Π represents swimming performance parameter, such as net force, power, cost of transport, etc.

**Fig 2.**
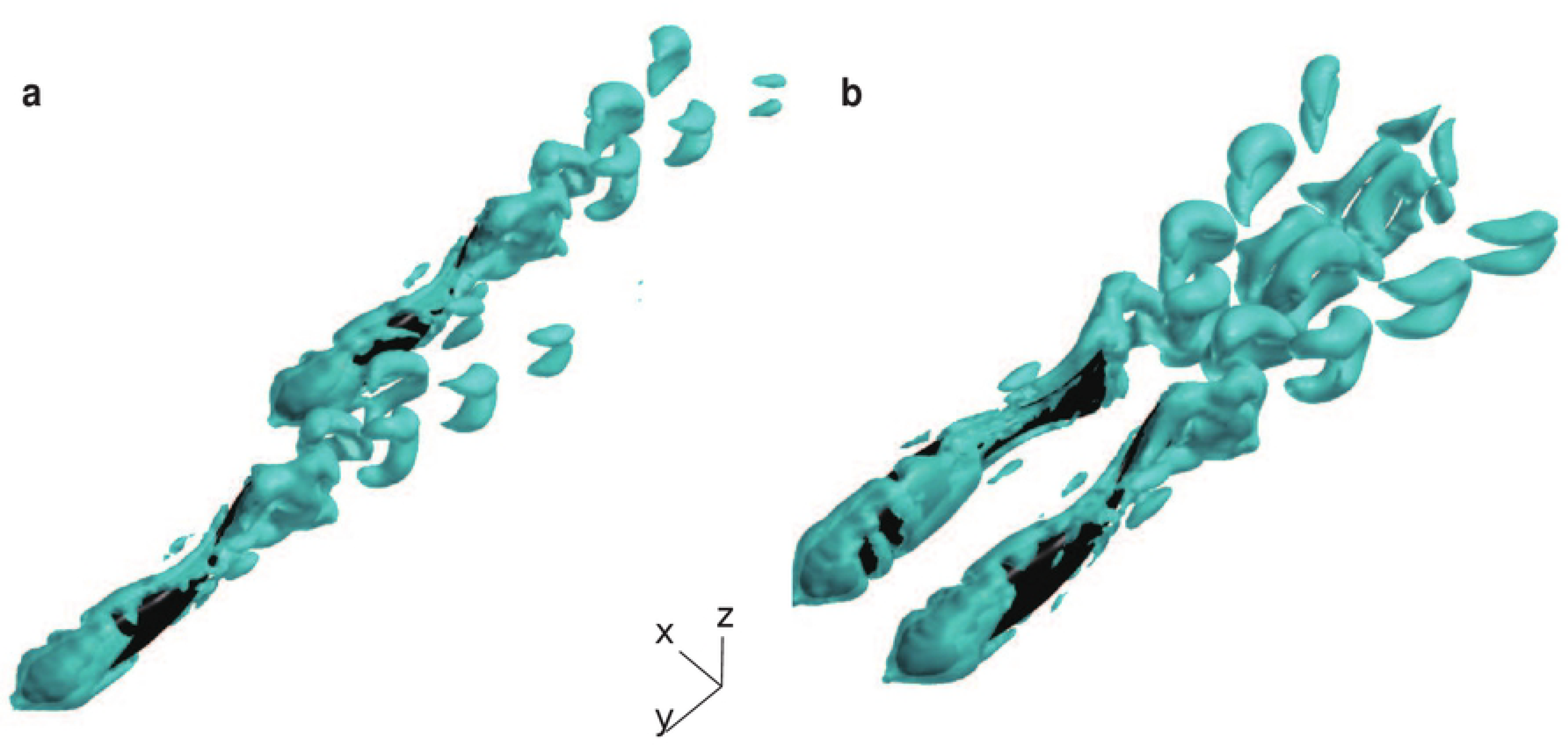
Three-dimensional flow visualization using an iso-surface of the Q-criterion [34]. (a) *δx* = 0.2*L*, *δy* = 1.25*L*, *δϕ* = 0; (b) *δx* = 0.5*L*, *δy* = 0, *δϕ* = *T /*2. For more examples of the wake topology, see [35].

## Results

We conducted a simulation of a solitary fish in self-propelled swimming mode and obtained its terminal speed of 9.25 cm s^*−*1^ with a tail beat frequency of 8 Hz, which agrees well with the experiments [28]. We then applied an oncoming uniform flow at that velocity *U* = 9.25 cm s^−1^ (which gives a Reynolds number *Re* = 3700) and the same tail beat frequency of *f* = 8 Hz for all the rest of simulations in tethered mode. Note that the speed and the kinematics are not chosen arbitrarily, but representatively: a range of speeds of approximately 3 to 15 cm s^−1^ has been observed in the experiments [28, Fig. 2], and 9.25 cm s^−1^ is almost in the middle. The experiments also suggested that fish had preferred combinations of frequency and amplitude depending on the speed. One of those is used in the simulations.

Thus, we obtained the swimming performance Π_*solo*_ of a solitary tethered fish and a performance map Π(*δx, δy, δϕ*) of the protagonist fish in pairwise simulations (see Fig. 1), where the symbol Π represents a time-average performance parameter such as net force, power, etc. Among a variety of performance parameters, we chose the net longitudinal force *F*_‖_ and hydrodynamic power *P* as indicators of propulsive efficiency, and the lateral force *F_⊥_*, standard deviation of longitudinal force *s.d.F*_‖_, and standard deviation of lateral force *s.d.F*_⊥_ as indicators of stability. We quantify the effect of hydrodynamic interaction either as residual difference ∆Π(*δx, δy, δϕ*) = Π(*δx, δy, δϕ*) − Π_*solo*_ or as percentage Π(*δx, δy, δϕ*)/Π_*solo*_ × 100%, whichever is more appropriate in its context.

### Effect on propulsive efficiency

#### Net longitudinal force

The hydrodynamic interaction between the two fish induces an extra longitudinal force ∆*F*_‖_ on the protagonist fish, which is shown in Fig. 3 in dimensionless form, normalized by the weight of the fish *mg*. The induced force ∆*F*_‖_ can act in the direction of drag or thrust, depending on the relative position of the two fish in the pair, and in magnitude it reaches 0.0018*mg*. Interestingly, ∆*F*_‖_ does not depend on *δϕ* as much as on *δx* and *δy*.

**Fig 3.**
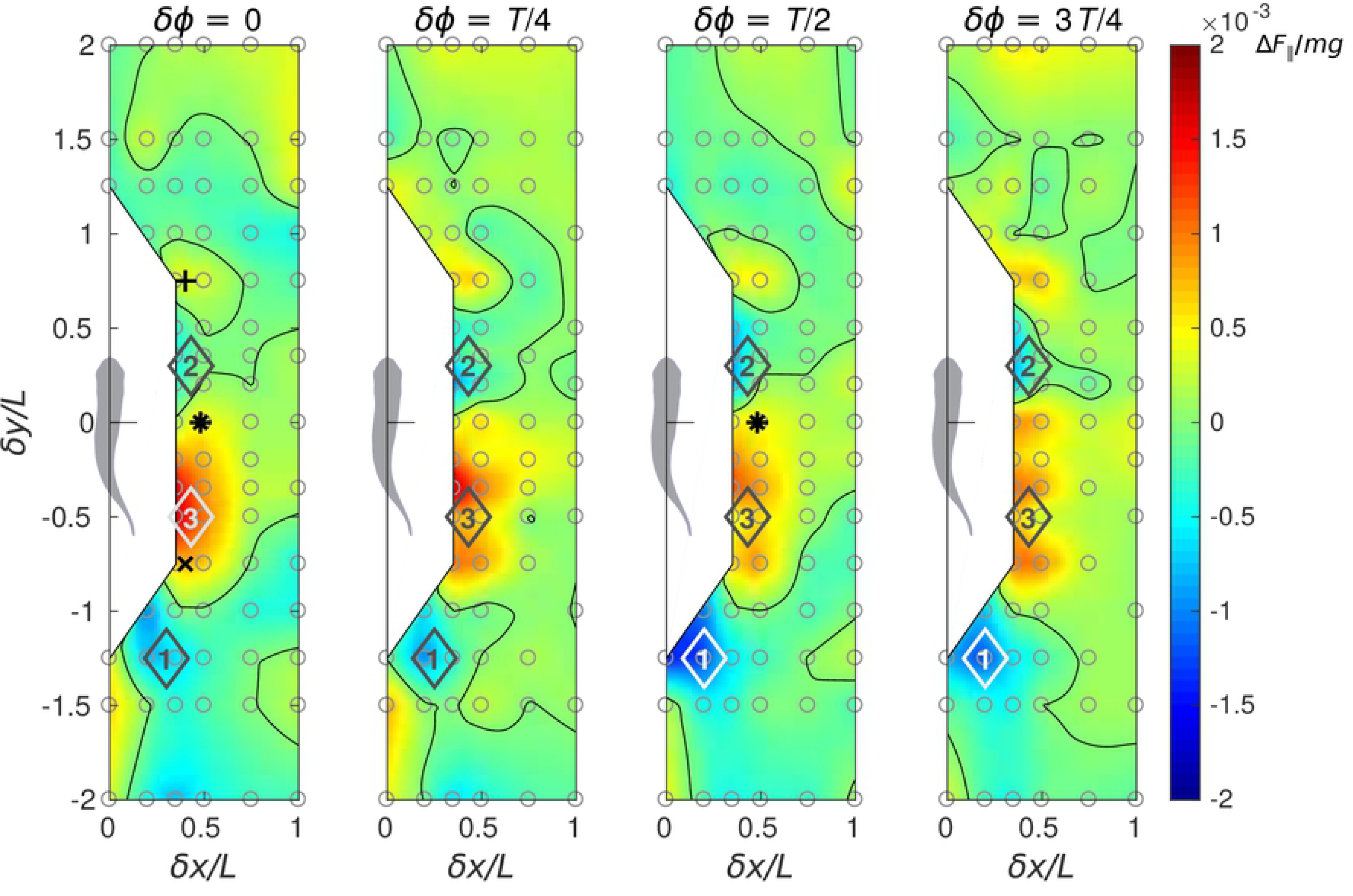
Performance maps of the protagonist fish in terms of normalized net longitudinal force, ∆*F*_‖_/*mg* = (*F*_‖_ − *F*_‖*solo*_)/*mg*.

The protagonist fish experiences the largest drag when it ineptly plunges into the wake of its companion. This regime corresponds to the blue spots situated between *δy* = −2*L* and *−L*, see locations 1 in Fig. 3, where *L* is the fish body length. Since these regions coincide with the positions of the vortices shed by the companion fish (see Fig. 2a), the increased drag can be regarded as a direct consequence of inflow turbulence. Upon receiving such a penalty in ∆*F*_‖_, the protagonist fish in free swimming would adopt a more powerful stroke to maintain its speed and position, otherwise it would decelerate and fall behind. A follower protagonist can experience thrust if placed in-line behind its leader companion, but this effect is confined to a narrow band (*δx <* 0.2*L*). The leader does not experience any substantial ∆*F*_‖_ when swimming in such tandem formation, i.e., there is no updraft. This finding contrasts with the strong upstream drafting observed in tandem arrangements of drag-generating flapping flags [33].

When the protagonist fish swims in a staggered side-by-side formation with its companion, in a slightly leading position (*δx <* 0.5*L* and 0 *< δy <* 0.6*L*, see locations 2 in Fig. 3) it experiences slightly negative ∆*F*_‖_, while in a slightly trailing position (*δx <* 0.7*L* and *−L < δy <* 0, see locations 3 in Fig. 3) the protagonist fish benefits from the largest positive ∆*F*_‖_. This implies that, in a staggered side-by-side formation, maximum extra thrust for the fish which is lagging behind entails drag for the fish which is leading. Therefore, in free swimming, this formation is likely to be unstable and to promote side-by-side arrangement with *δy* ≈ 0 (as in Fig. 2b) so that the two fish equalize. Earlier experiments [28] using red nose tetra fish indeed showed that a pair of fish preferred phalanx formations with *δx ≈* 0.6*L* and −0.2*L < δy <* 0.

#### Power

The hydrodynamic power consumption *P* of one fish in a pair varies with *δx*, *δy* and *δϕ* and it generally differs by less than 10% from *P*_*solo*_, see Fig. 4. Remarkably, swimming with *δϕ* = 0 and *T*/2 is substantially less demanding in terms of power requirements than with *δϕ* = *T*/4 or 3*T*/4 (by up to approximately 5%). This may explain the preference for either in-phase or anti-phase synchronization observed in tetra fish in high energy demanding swimming regimes [28].

**Fig 4.**
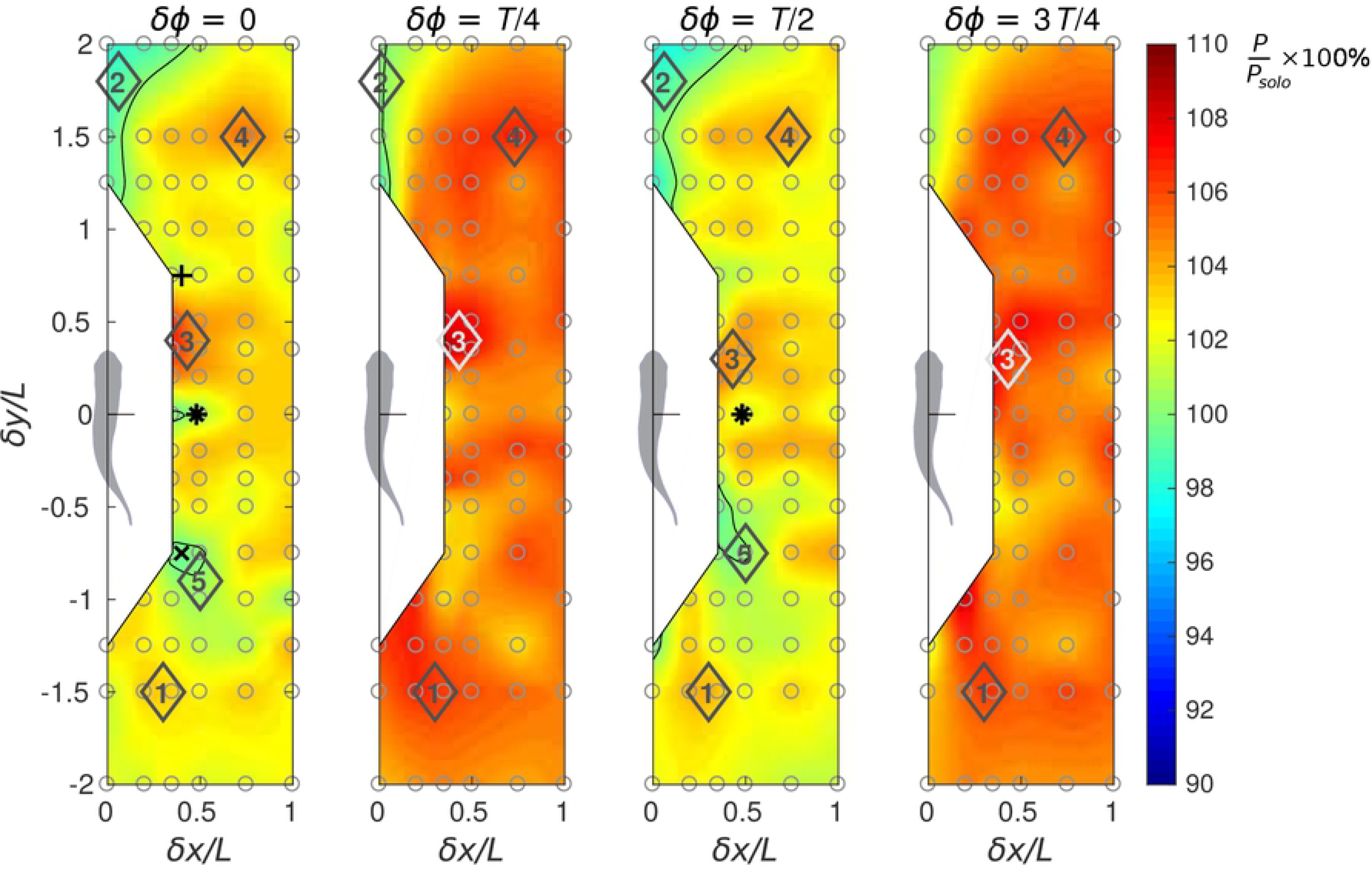
Performance maps of the protagonist fish in terms of relative power consumption, *P*/*P*_*solo*_ × 100%.

Relative spatial positioning also matters. When a follower protagonist fish is swimming in the vortex wake behind its leader companion, in addition to the increased drag, it spends more power (locations 1, Fig. 4). In this formation, the follower’s power consumption rises up to *P/P*_*solo*_ = 1.03 if the phasing is favorable (*δϕ* = 0) and to *P/P*_*solo*_ = 1.08 if the phasing is unfavorable (*δϕ* = *T /*4). Hence, this relative position must be avoided for being energetically inefficient. Positioning straight ahead of the companion fish may slightly lower the power consumption (locations 2, Fig. 4).

Swimming on a diagonal in front of the companion fish may increase the power consumption (locations 3 and 4 in Fig. 4), while swimming on a diagonal behind the companion (locations 5 in Fig. 4) requires less power if *δϕ* = 0 or *T /*2. Staggered side-by-side formations are appealing when both the follower and the leader can enjoy extra thrust at negligible energetic cost. For instance, the case *δϕ* = 0, *δx* = 0.4*L* and *δy* = ∓0.75*L* (‘+’ and ‘*×*’ symbols in Figs. 3 and 4) shows ∆*F*_‖_/*mg* = 0.0004, *P*/*P*_*solo*_ = 1 for the follower and ∆*F*_‖_/*mg* = 0.0007, *P*/*P*_*solo*_ = 1.01 for the leader, respectively. This means net benefit for the follower and possible benefit for the leader since, by a slight decrease in tail beat amplitude, thrust can be converted into power savings. However, swimming in this staggered formation requires good control skill from both individuals to keep the overall favorable relative position.

A side-by-side formation with *δx* = 0.48*L* and *δy* = 0 (‘*\**’ symbols in Figs. 3 and 4) yields ∆*F*_‖_/*mg* = 0.0004, *P*/*P*_*solo*_ = 1 for both individuals if they are synchronized in-phase, and ∆*F*_‖_/*mg* = 0.0005, *P*/*P*_*solo*_ = 1.01 for both if they swim in anti-phase. This condition may be acceptable for the fish from the energetic point of view.

#### A comprehensive result by *CoT* accounting for gait adjustment

Fish in schooling configurations need to adjust their gait to maintain their relative position. In our numerical simulations, we have prescribed the same tail beat frequency *f* and midline deformation envelope with amplitude *a* for both fish in the pair, see Methods. Real fish may adjust these parameters to reach the objective of steady swimming, but to remain synchronized, the group members must maintain equal frequency, while midline deformation can be used as a free control parameter.

The hydrodynamic interactions are weak enough to estimate the necessary adjustment of *a* using linear extrapolation. We therefore carry out an additional solitary fish simulation with *a* increased by 5%, i.e, 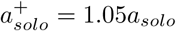, where the subscript ‘*solo*’ stands for the solitary fish. We use ‘+’ when we refer to the results of this additional simulation, and no superscript for the original simulation. The derivative of the longitudinal force with respect to the amplitude and the derivative of the power with respect to the longitudinal force are approximated as, respectively,

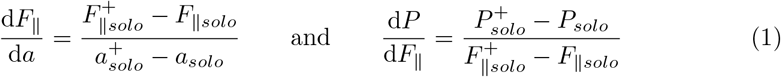

at *a* = *a*_*solo*_. The above derivatives are used for calculating the adjusted amplitude 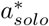 and power 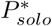 of the solitary fish that would correspond to steady swimming at the same prescribed velocity *U*,

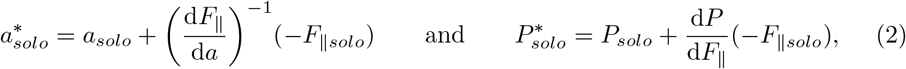

by ensuring the longitudinal force be close to 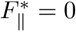. Similarly, for all points on the two-fish school diagrams versus separation *δx* and *δy* between the fish, we calculate

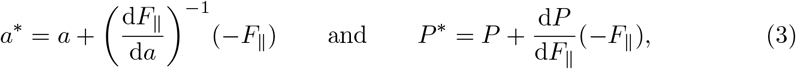

In (3), the values of *a*, *P* and *F*_‖_ correspond to the simulation data for the protagonist fish in the pair, for which the diagram is made.

There exist several different criteria commonly used to evaluate energetic efficiency of self-propelled swimming [36]. In this study, we choose the cost of transport *CoT* for its intuitive physical interpretation as energy consumed per distance traveled, which after normalization by the body weight becomes equal to

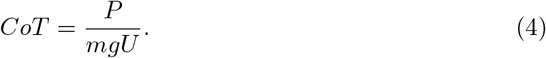

Note that a direct application of this formula to the results of our numerical simulations would be problematic, because (4) implies that the fish is in steady forward swimming, which is in contradiction to the non-zero net longitudinal force in the simulations (see Fig. 3). This problem is solved by using extrapolation to estimate the power under zero-longitudinal-force condition, as explained above.

The values of *P*^*^ and 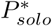 determined from (3) and (2) all correspond to the same swimming speed *U*. The cost of transport is thus equal to *P*^*^/*mgU* and 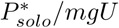, respectively. Therefore, the energetic benefit for the second fish in a pair can be quantified using the *CoT* ratio

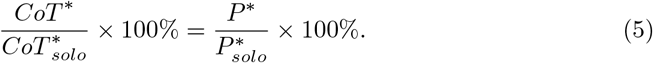

It should be reminded that our estimate is based on a linear approximation, i.e., all quadratic and higher order terms are neglected in (3). Therefore, if *a*^*^ differs from *a* by about 10%, one can expect an order of magnitude of 1% for the approximation error. In addition, (3) only corrects for the longitudinal force, but the lateral force remains unbalanced. Finally, the estimate includes the numerical simulation error due to the limitation of fixed CoM, absence of control, etc.

As shown in Fig. 5, when the phase difference between the two fish is *δϕ* = *T /*4 or 3*T /*4, for the protagonist fish, the estimated cost of transport is globally greater than that of the in-phase and antiphase (*δϕ* = 0 or *T /*2). For all phase shift conditions, when the protagonist fish is exposed to the wake (vortex street) of the companion fish (locations 1, Fig. 5), greater cost of transport is incurred. If the protagonist fish is located ahead of the companion fish (locations 2, Fig. 5), it may slightly decrease the cost of transport. However, in that condition, the companion fish is located in the wake of the protagonist fish and it may prefer to relocate. In a staggered side-by-side formation, if the protagonist fish is slightly in front (locations 3, Fig. 5), strong negative interaction occurs. Contrarily, if the protagonist fish is slightly behind, it can receive energetic benefit (locations 4). Still, in this situation, the companion fish is located on the diagonal in front of the protagonist fish and experiences negative influence. The relative positions of the fish pairs studied experimentally by Ashraf et al. [28] are indicated by triangles in Fig. 5. The fish appear to avoid the regions of strong variation in the cost of transport.

**Fig 5.**
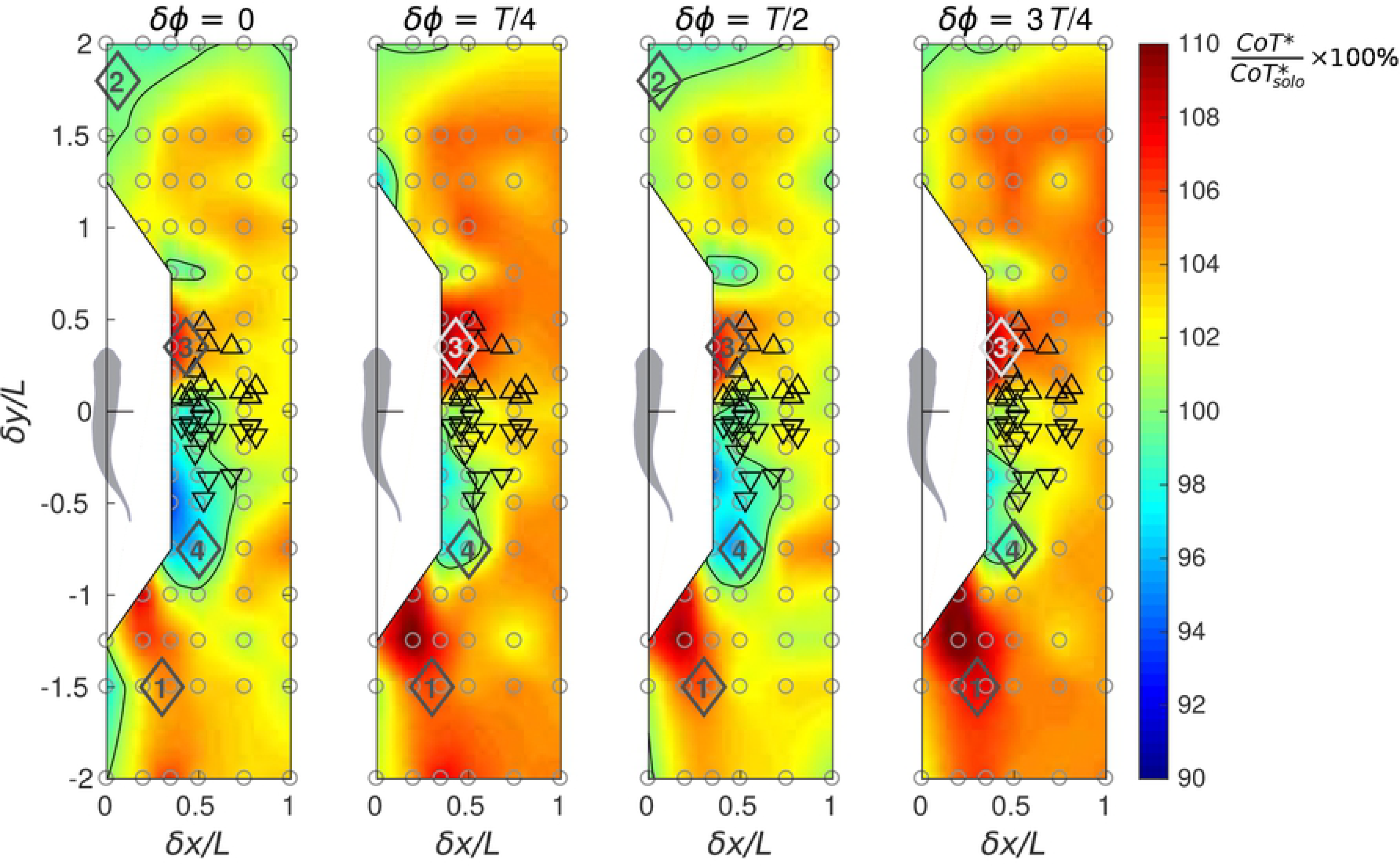
Performance maps of the protagonist fish in terms of cost of transport, 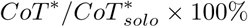.

### Effect on stability

To study the effect of schooling on the stability of the fish pair swimming pattern, we examine the lateral forces and the fluctuation (represented by the standard deviation) of lateral and longitudinal forces. These results for in-phase swimming are summarized in the diagrams in Fig. 6.

**Fig 6.**
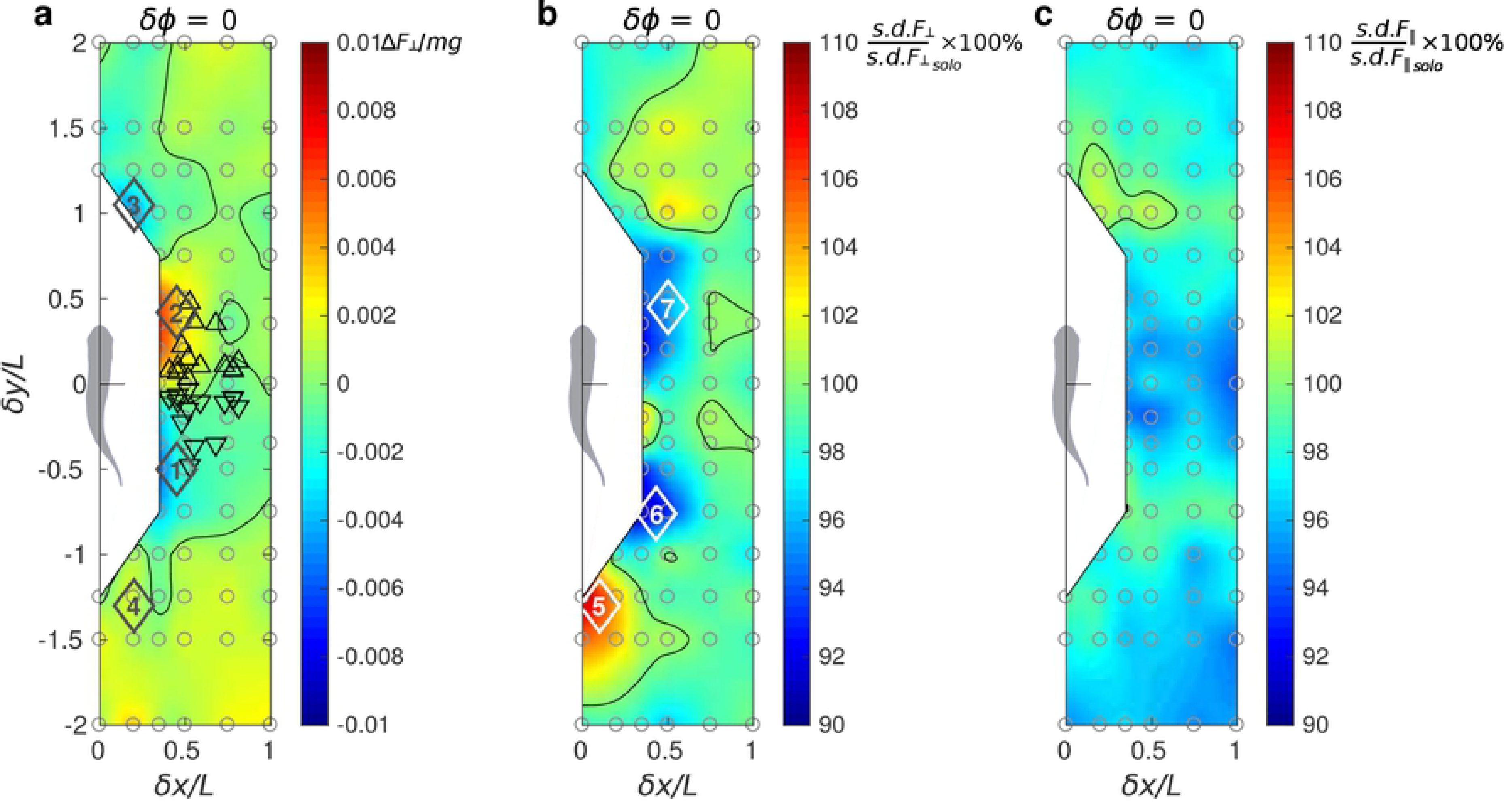
Performance maps of the protagonist fish in terms of (*a*) nomalized lateral force, ∆*F*_⊥_/*mg*, (*b*) relative standard deviation of lateral force, *s.d.F*_⊥_/*s.d.F*_⊥*solo*_ × 100% and (*c*) relative standard deviation of longitudinal force, *s.d.F*_‖_/*s.d.F*_‖solo_ × 100%.

#### Lateral force

Receiving unbalanced lateral force may break the stable configuration between the two fish, unless the fish spends more effort to adjust the unbalanced lateral force to maintain their relative position, but such effort may reduce the energetic efficiency. In our numerical simulations, the fish does not implement such adjustment, since we prescribe a bilaterally symmetric body deformation envelop, see Material and Methods. Instead, the fish is free to rotate about its CoM. While the time-average body orientation remains precisely forward in solitary swimming, it becomes significantly deflected to the left or to the right as soon as the flow symmetry is broken by the presence of a companion fish. Therefore, *F*_⊥_ includes contributions from two hydrodynamic interaction effects: bilateral asymmetry in the surface pressure distribution and reorientation of the fish in the laboratory reference frame.

As shown in Fig. 6a, there are several locations that could lead to dramatically unbalanced lateral force: when the two fish swim side by side, a slight trailing position (location 1) pulls the protagonist fish towards its companion, on the contrary, a slight leading position pushes it apart (location 2). The two fish in a side-by-side configuration may align themselves (*δy* ≈ 0) to keep away from the zones of strong unbalance, which seems to agree with the behaviors in the experiments [28] (triangles in Fig. 6a). Also, leading (location 3) and trailing (location 4) positions may also produce lateral imbalance. The results shown in Fig. 6a for in-phase swimming are representative of all synchronizations.

#### Fluctuation of force

Within one tail beat cycle, the force exerted on the fish body fluctuates quasi-periodically. The fluctuation of lateral and longitudinal forces may also affect the stability in fish swimming. Halsey et al. [14] notice that fish may not be able to maintain station relative to their neighbors when they swim in a turbulent water stream. It is logical to conjecture that the leader’s wake can have a similar impact on the followers even if the ambient flow is laminar. Here, we utilize the standard deviation of the lateral and the longitudinal forces to quantify the fluctuation. Figures 6b and c show, respectively, *s.d.F*_⊥_ and *s.d.F*_‖_ for the in-phase synchronized cases. When comparing between these two components, it is important to bear in mind that, for a solitary swimmer, the lateral force fluctuation is three times as strong as the longitudinal force fluctuation, i.e., *s.d.F*_⊥solo_/*mg* = 0.0046 while *s.d.F*_*‖solo*_/*mg* = 0.0016. Figs. 6b and c only show how this fluctuation is amplified of attenuated due to hydrodynamic interaction between the two fish when they swim in a pair. Thus, *s.d.F*_⊥_ can differ from *s.d.F*_⊥*solo*_ by as mush as ±9%, while *s.d.F*_‖_ only differs from *s.d.F*_*‖solo*_ by between −6% and +2%. These facts taken together, we conclude that fluctuation in the lateral direction is more likely to be a strong destabilizing factor. This situation also holds for *δϕ* = *T /*4, *T /*2 and 3*T /*4 (not shown). Considering the spatial structure of *s.d.F*_⊥_ when *δϕ* = 0 (Fig. 6b), we notice that location 5 corresponds to strengthened fluctuation that the fish may avoid. The staggered side-by-side locations 6 and 7 may be chosen to attenuate the lateral fluctuation. It should be mentioned, however, that the spatial position of the peaks of *s.d.F*_⊥_ varies with *δϕ* (not shown).

## Discussion

Our results show that the spatial organization and the kinematic synchronization of a pair of swimming fish—the *minimal school*—have a clear effect on two crucial aspects of schooling: energy expenditure and fluctuation minimization. We have examined the effect of the hydrodynamic interaction between the two fish on several performance parameters by probing forces and consumed hydrodynamic power on a fish that we have called the protagonist fish, while placing it in different positions and with a kinematic phase shift with respect to its neighbor (the companion fish).

Regarding energy expenditure, we have used a cost of transport function (see Fig. 5) that brings out two main conclusions. On the one hand, swimming in phase (*δϕ* = 0) or anti-phase (*δϕ* = *T/*2) is advantageous over the cases of quarter-period phase shift (*δϕ* = *T/*4 and 3*T/*4). Yet, it remains to be clarified whether the prevalence of in- and anti-phase lock behavior [28] stems from mechanical coupling akin to flagellar synchronization [37, 38] or from sensorimotor abilities. On the other hand, regardless of the phasing between neighbors, certain relative positions are beneficial or penalizing. Most notably, a side-by-side configuration with the protagonist fish slightly diagonally behind is beneficial for the protagonist fish, while lagging behind in the region of the wake of the companion fish is penalizing. When comparing the cost of transport maps with the positions of an experiment with a pair of tetra fish (triangles in Fig. 5), high cost of transport zones appear to be avoided by the fish.

Considering the mechanisms such as updraft and channeling effect, as the number of fish involved in the collective behavior increases, the hydrodynamic benefit may accumulate as a quasi-steady linear interaction. These long-range interactions may be described analytically using dipolar far-field approximation [39]. Conversely, our results suggest that, as the number of fish decreases to two, unsteady and nonlinear interaction between the two fish becomes non negligible and specific flow structures and phase differences become important factors. It remains to be investigated how hydrodynamic influence evolves as the number of fish in a school increases.

Concerning the wake energy harvesting mechanism, our results suggest that, when a fish locates in the wake of the upfront leading fish, it becomes energetically inefficient. However, one should be aware that our conclusion is based on the tethered motion (fixed CoM) and absence of kinematic adjustment. A recent study by Verma et al. [27] shows that, when learning-based optimized kinematic adjustment is present, wake capture can be advantegeous. Therefore, the comparison between the present study and study of Verma et al. [27] demonstrates that there exists a distinction between wake capturing and wake energy harvesting: successful wake capture requires skills in sensing and adjustment, and if the fish (or an artificial swimmer) lacks those skills, wake capture may become energetically unfavorable. Besides the active mechanism, passive mechanisms based on appropriate body flexibility and mass distribution are also potential factors that may influence fish performance in school [25].

We hypothesize that fish avoid wake capturing and adopt side-to-side configuration as a conservative strategy when energy harvesting is impractical due to adverse environmental conditions, physiological constraints, or other impeding factors. Furthermore, in comparison with two-dimensional wakes, three-dimensional fish wakes are geometrically more complex and less stable. The energy of vortex motion rapidly cascades to small-scale structures and dissipates, which hinders wake energy harvesting in 3D (cf. 36% decrement of *CoT* in 2D and only 5% decrement in 3D, in Verma et al. [27]). Further study is needed to quantify and fundamentally explain the difference between hydrodynamic interactions in the two-dimensional and the three-dimensional contexts.

The stability of the school has been studied examining the lateral forces and fluctuations of both lateral and longitudinal forces as functions of the relative position and kinematic phase shift. All phase lags produce qualitatively the same picture concerning lateral force and fluctuations, hence Fig. 6 where only the *δϕ* = 0 case is shown. The fish would need to compromise between propulsive efficiency and stability, since the optimal positions for *CoT* and lateral force fluctuations do not coincide (for instance, compare between Fig. 5a and Fig. 6a). Fish seeking for a stable position may suffer from high *CoT* and *vice versa*. In addition, fish may seek for mutually beneficial formations, since a schooling configuration exclusively beneficial to one member may be severely unfavorable to the other. A stable fish school configuration ought to be a concord between all members.

## Methods

We developed an in-house three-dimensional overset grid numerical approach based on finite-volume method and programmed in FORTRAN 90 to simulate cyclic swimming of fish [29–31, 40]. The approach comprises surface models of the changing fish shape (dimension: 121 × 97), and local fine-scale body-fitted grids (dimension: 121 × 97 × 20) plus a large stationary global grid (dimension: various) to calculate the flow patterns around the fish with sufficient resolution (supportive information on grid resolution and size tests can be found in supplementary materials). As shown in Figure 7, to simulate a fish pair, two body-fitted grids were deployed, which deformed as the fish model deformed. The global grid surrounded the body-fitted grids and covered a sufficiently large domain to enclose the swimming fish and their wake. The ensemble of body-fitted grids and global grid was set up as a multi-blocked, overset-grid system based on a chimera grid scheme [41]. During the simulation, the body-fitted and global grids share values on their interfaces through inter-grid communication algorithm. The body was modelled on the silhouette of a Red nose tetra fish (*Hemigrammus bleheri*), with a body length of 4 cm, an average length measured in previous experimental study [28]. All cross-sections of the fish were modeled as ellipses. To reduce the complexity in modeling and computation, we assume that the hydrodynamic influence of all fins other than the tail fin is relatively minor, and neglect them in the model. Also, for the same reason, the gap of the fork-shaped tail fin was neglected, and the fish model has a triangle-shaped fin instead. The instantaneous body shape was driven by:

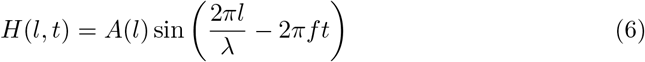

where *l* is the dimensionless distance from the snout along the longitudinal axis of the fish based on the length of the fish model *L*; *H*(*l, t*) is the dimensionless lateral excursion at time *t*;

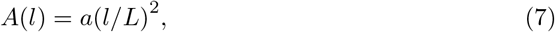

is the dimensionless amplitude envelope function at *l*; *λ* is the length of the body wave and it is set as 1.2*L*; *f* is the tail beat frequency defined as *f* =8 Hz, a typical value in the experimentals [28]. We use *a* = 0.11 in all simulations, unless stated. Eq. 6 may cause total body length along the midline to vary during the tail beat; this variation is corrected by a procedure that preserves the lateral excursion *H*(*l, t*) while ensuring that the body length remains constant. Procedure flow of simulations is shown in Fig. 1*b*. We conducted simulations in two modes. In free-swimming (self-propelled) mode simulation, we simulated single fish swims in the horizontal plane with its center-of-mass (CoM) movements and body orientation determined by the hydrodynamic forces on the body, while oncoming flow was set as zero. By using free-swimming simulation, we obtained the terminal speed in single fish swimming and apply to the rest simulations. All the rest simulations were conducted in fixed CoM mode: we simulated a single fish or fish pair swimming with CoM and relative position fixed, while the rotational degree of freedom was still available to model the rotational recoil effect during swimming. The oncoming flow was set as the terminal speed obtained in free-swimming simulation. The Reynolds number of the simulations is defined as *Re* = *ρU L/µ*, where *ρ* is the water density, *U* is the swimming speed, *L* is the body length, and *µ* is the dynamic viscosity of water. The free-swimming simulation on a single fish rendered an equilibrium speed of 9.25 cm s^−1^. In all the rest simulations, the Re was set as 3700, and no turbulence model was applied in the simulation.

**Fig 7.**
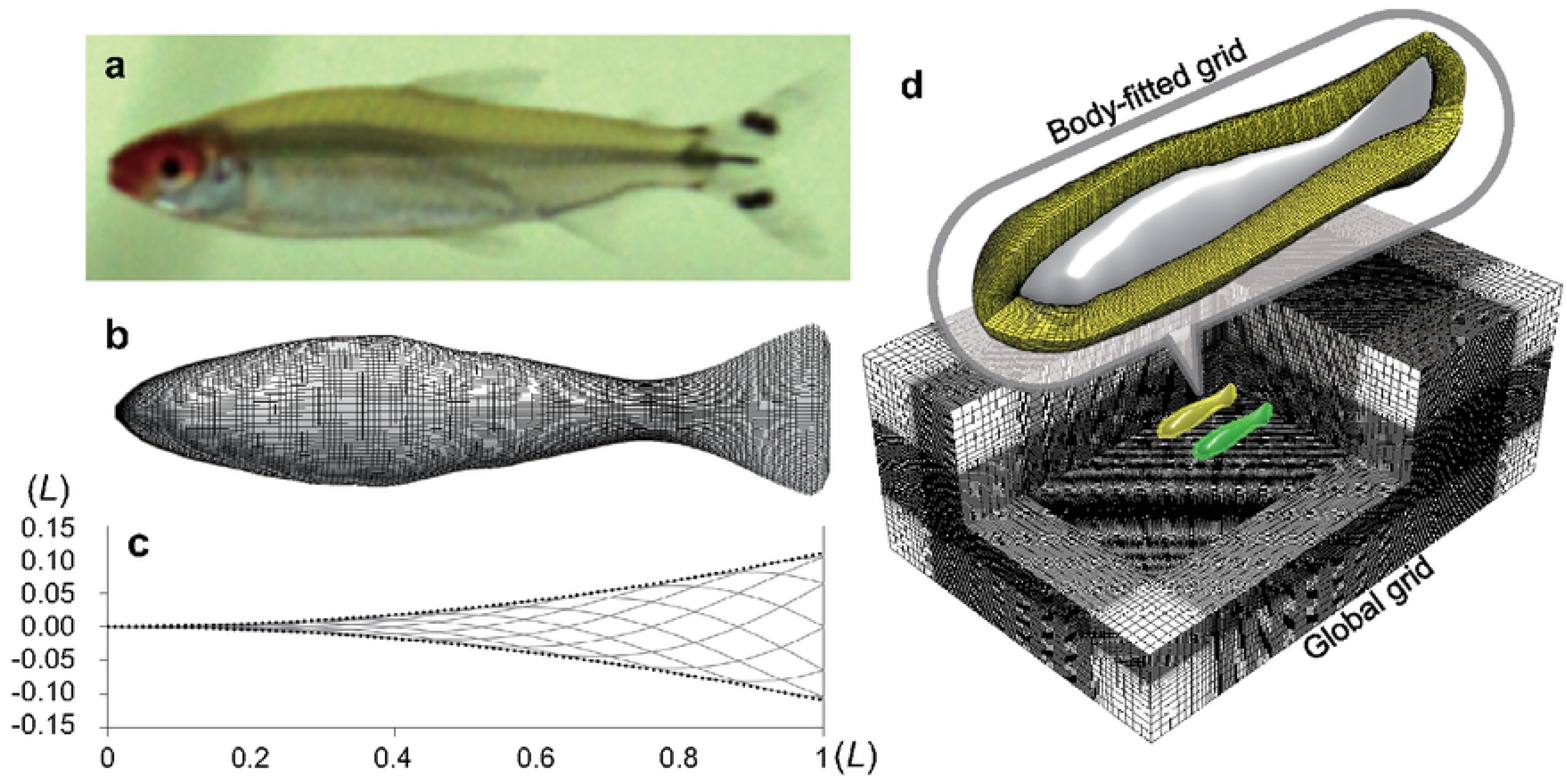
(a) Red nose tetra fish (*Hemigrammus bleheri*); (b) Surface model of Red nose tetra fish (dimension: 121 × 97); (c) A function (eq. 6) drives the instantaneous body shape. Variation of body length caused by this driving function was corrected to keep lateral excursion and body length constant at 1*L*. (d) Multi-blocked computational grid system consists of local fine-scale body-fitted grids (dimension: 121 × 97 × 20) plus a large stationary global grid (dimension: various).

The simulations on a fish pair were implemented by varying the relative longitudinal and lateral positions between two fish. The choice of using fixed CoMs for the two fish ensured that the relative position between the fish during swimming was unaffected by their complex interaction. Meanwhile, to test the influence of phase difference, for each position we implemented four simulations with varied phase shift between the two fish (*δϕ* = 0, *T/*4, *T/*2, 3*T/*4, respectively). Based on 312 of simulation results and interpolation among those results, we could construct the swimming performance map for different performance parameters. Fig. 1*a* explains how to comprehend the performance maps (Figs. 3-6). Note that a performance map is not a result of one simulation, but a summation of many simulations with a same phase shift between the two fish. Each circle in the map represents a simulated case, and the value at this point is the swimming performance (force, power, etc.) of the protagonist fish. The companion fish is placed at the origin point, while protagonist fish is assumed to be deployed in a range of relative position, which covers ±2*L* in longitudinal direction and from 0 to 1*L* in the lateral direction. Our definition of power is explained in the Electronic Supplementary Material. We calculate the time variation of the hydrodynamic power and apply time-averaging over one tail beat cycle.

## Acknowledgements

G.L. is funded by the Japan Society for the Promotion of Science (JP17K17641), and the Sasakawa Scientific Research Grant 2018-7022 from The Japan Science Society. D.K. is supported by the Japan Society for the Promotion of Science (JP18K13693). H.L. is partly supported by the Grant-in-Aid for Scientific Research on Innovative Areas of No. 24120007, JSPS.

## References

1. Czirók A, Vicsek M, Vicsek T. Collective motion of organisms in three dimensions. Physica A: Statistical Mechanics and its Applications. 1999;264(1):299–304. doi:10.1016/S0378-4371(98)00468-3.

2. Vicsek T, Zafeiris A. Collective motion. Physics Reports. 2012;517(3):71–140. doi:10.1016/j.physrep.2012.03.004.

3. Shaw E. Schooling fishes. Scientific American. 1978;66(2):166–175.

4. Partridge BL. The structure and function of fish schools. Scientific American. 1982;246(6):114–123.

5. Lopez U, Gautrais J, Couzin ID, Theraulaz G. From behavioural analyses to models of collective motion in fish schools. Interface Focus. 2012;2(6):693–707. doi:10.1098/rsfs.2012.0033.

6. Partridge BL, Pitcher TJ. The sensory basis of fish schools: Relative roles of lateral line and vision. Journal of comparative physiology. 1980;135(4):315–325. doi:10.1007/BF00657647.

7. Breder CM. Vortices and fish schools. Zoologica: scientific contributions of the New York Zoological Society. 1965;50:97–114.

8. Liao JC. A review of fish swimming mechanics and behaviour in altered flows. Philosophical Transactions of the Royal Society B: Biological Sciences. 2007;362(1487):1973–1993. doi:10.1098/rstb.2007.2082.

9. Weihs D. Hydromechanics of Fish Schooling. Nature. 1973;241:290–291.

10. Fields PA. Decreased swimming effort in groups of pacific mackerel (*Scomber japonicus*). American Zoologist. 1990;30(4):A134–A134.

11. Herskin J, Steffensen JF. Energy savings in sea bass swimming in a school: measurements of tail beat frequency and oxygen consumption at different swimming speeds. Journal of Fish Biology. 1998;53(2):366–376. doi:10.1111/j.1095-8649.1998.tb00986.x.

12. Johansen JL, Vaknin R, Steffensen JF, Domenici P. Kinematics and energetic benefits of schooling in the labriform fish, striped surfperch *Embiotoca lateralis*. Marine Ecology Progress Series. 2010;420:221–229. doi:10.3354/meps08885.

13. Marras S, Killen SS, Lindström J, McKenzie DJ, Steffensen JF, Domenici P. Fish swimming in schools save energy regardless of their spatial position. Behavioral Ecology and Sociobiology. 2015;69(2):219–226. doi:10.1007/s00265-014-1834-4.

14. Halsey LG, Wright S, Racz A, Metcalfe JD, Killen SS. How does school size affect tail beat frequency in turbulent water? Comparative Biochemistry and Physiology Part A: Molecular & Integrative Physiology. 2018;218:63–69. doi:10.1016/j.cbpa.2018.01.015.

15. Ashraf I, Bradshaw H, Ha TT, Halloy J, Godoy-Diana R, Thiria B. Simple phalanx pattern leads to energy saving in cohesive fish schooling. Proceedings of the National Academy of Sciences. 2017;114(36):9599–9604. doi:10.1073/pnas.1706503114.

16. Bergmann M, Iollo A. Modeling and simulation of fish-like swimming. Journal of Computational Physics. 2011;230(2):329–348. doi:10.1016/j.jcp.2010.09.017.

17. Gazzola M, Chatelain P, van Rees WM, Koumoutsakos P. Simulations of single and multiple swimmers with non-divergence free deforming geometries. Journal of Computational Physics. 2011;230(19):7093–7114. doi:10.1016/j.jcp.2011.04.025.

18. Gazzola M, Hejazialhosseini B, Koumoutsakos P. Reinforcement learning and wavelet adapted vortex methods for simulations of self-propelled swimmers. SIAM Journal on Scientific Computing. 2014;36(3):B622–B639. doi:10.1137/130943078.

19. Hemelrijk CK, Reid DAP, Hildenbrandt H, Padding JT. The increased efficiency of fish swimming in a school. Fish and Fisheries. 2015;16(3):511–521. doi:10.1111/faf.12072.

20. Becker AD, Masoud H, Newbolt JW, Shelley M, Ristroph L. Hydrodynamic schooling of flapping swimmers. Nature Communications. 2015;6:8514. doi:10.1038/ncomms9514.

21. Novati G, Verma S, Alexeev D, Rossinelli D, van Rees WM, Koumoutsakos P. Synchronisation through learning for two self-propelled swimmers. Bioinspiration & Biomimetics. 2017;12(3):036001.

22. Maertens AP, Gao A, Triantafyllou MS. Optimal undulatory swimming for a single fish-like body and for a pair of interacting swimmers. Journal of Fluid Mechanics. 2017;813:301–345. doi:10.1017/jfm.2016.845.

23. Park SG, Sung HJ. Hydrodynamics of flexible fins propelled in tandem, diagonal, triangular and diamond configurations. Journal of Fluid Mechanics. 2018;840:154–189. doi:10.1017/jfm.2018.64.

24. Peng ZR, Huang H, Lu XY. Collective locomotion of two closely spaced self-propelled flapping plates. Journal of Fluid Mechanics. 2018;849:1068–1095. doi:10.1017/jfm.2018.447.

25. Dai L, He G, Zhang X, Zhang X. Stable formations of self-propelled fish-like swimmers induced by hydrodynamic interactions. Journal of The Royal Society Interface. 2018;15(147). doi:10.1098/rsif.2018.0490.

26. Daghooghi M, Borazjani I. The hydrodynamic advantages of synchronized swimming in a rectangular pattern. Bioinspiration & Biomimetics. 2015;10(5):056018.

27. Verma S, Novati G, Koumoutsakos P. Efficient collective swimming by harnessing vortices through deep reinforcement learning. Proceedings of the National Academy of Sciences. 2018;115(23):5849–5854. doi:10.1073/pnas.1800923115.

28. Ashraf I, Godoy-Diana R, Halloy J, Collignon B, Thiria B. Synchronization and collective swimming patterns in fish (Hemigrammus bleheri). Journal of the Royal Society Interface. 2016;13(123). doi:10.1098/rsif.2016.0734.

29. Li G, Müller UK, van Leeuwen JL, Liu H. Body dynamics and hydrodynamics of swimming fish larvae: a computational study. Journal of Experimental Biology. 2012;215(22):4015–4033. doi:10.1242/jeb.071837.

30. Li G, Müller UK, van Leeuwen JL, Liu H. Escape trajectories are deflected when fish larvae intercept their own C-start wake. Journal of The Royal Society Interface. 2014;11(101). doi:10.1098/rsif.2014.0848.

31. Li G, Müller UK, van Leeuwen JL, Liu H. Fish larvae exploit edge vortices along their dorsal and ventral fin folds to propel themselves. Journal of The Royal Society Interface. 2016;13(116). doi:10.1098/rsif.2016.0068.

32. Ashraf I, Godoy-Diana R, Halloy J, Collignon B, Thiria B. Supplementary material from “Synchronization and collective swimming patterns in fish Hemigrammus bleheri”; 2016. Available from: https://rs.figshare.com/collections/Supplementary_material_from_Synchronization_and_collective_swimming_patterns_in_fish_i_Hemigrammus_bleheri_i_/3500367/1.

33. Ristroph L, Zhang J. Anomalous hydrodynamic drafting of interacting flapping flags. Physical Review Letters. 2008;101:194502. doi:10.1103/PhysRevLett.101.194502.

34. Hunt JCR, Wray AA, Moin P. Eddies, streams, and convergence zones in turbulent flows. In: Studying Turbulence Using Numerical Simulation Databases. vol. 2; 1988.

35. Li G, Kolomenskiy D, Liu H, Thiria B, Godoy-Diana R. On the interference of vorticity and pressure fields of a minimal fish school. Submitted manuscript. Journal of Aero Aqua Bio-mechanisms. 2019;8(1).

36. Li G, Liu H, Muller UK, Voesenek CJ, van Leeuwen JL. Hydrodynamic cost of transport determines optimisation strategy: larval fish provide insight into undulatory swimming. Submitted manuscript. 2019.

37. Brumley DR, Wan KY, Polin M, Goldstein RE. Flagellar synchronization through direct hydrodynamic interactions. eLife. 2014;3:e02750. doi:10.7554/eLife.02750.

38. Kawamura Y, Tsubaki R. Phase reduction approach to elastohydrodynamic synchronization of beating flagella. Phys Rev E. 2018;97:022212. doi:10.1103/PhysRevE.97.022212.

39. Filella A, Nadal F, Sire C, Kanso E, Eloy C. Model of collective fish behavior with hydrodynamic interactions. Physical Review Letters. 2018;120:198101. doi:10.1103/PhysRevLett.120.198101.

40. Liu H. Integrated modeling of insect flight: from morphology, kinematics to aerodynamics. Journal of Computational Physics. 2009;228(2):439–459. doi:10.1016/j.jcp.2008.09.020.

41. Prewitt NC, Belk DM, Shyy W. Parallel computing of overset grids for aerodynamic problems with moving objects. Progress in Aerospace Sciences. 2000;36(2):117–172. doi:10.1016/S0376-0421(99)00013-5.

